# The lysosomal cation channel TRPML1 regulates the oligodendrocyte cytoskeleton

**DOI:** 10.64898/2026.06.16.731945

**Authors:** Lindsay K. Festa, Nikole Fandino Pachon, Ryan N. Anderson, Serena J. Chen, Alec L. Sciutto, Judith B. Grinspan, Kelly L. Jordan-Sciutto

## Abstract

Differentiating oligodendrocytes undergo dramatic morphologic alterations to transition from progenitors to mature oligodendrocytes that synthesize myelin, the lipid-rich membrane coating axons which strengthens saltatory conduction and provides metabolic support. Actin dynamics, which are often regulated by membrane bound nucleators associated with organelles, underpin the morphologic shifts in oligodendrocyte maturation; however, the origin of such regulation during oligodendrocyte differentiation remains unknown. Here, we demonstrate that the lysosomal non-selective cation channel, transient potential mucolipin 1 (TRPML1), is a critical regulator of oligodendrocyte morphology during differentiation and initial myelination. Lysosomes move into oligodendrocyte processes during differentiation. While manipulation of TRPML1 did not change the expression of oligodendrocyte lineage markers, activation of TRPML1 resulted in altered oligodendrocyte morphology and an increase in actin filament content driven by the small GTPase Rac1 and subsequent disinhibition of PAK1 via phosphorylation. Actin associated changes in morphology are accompanied by the presence of lysosomal-derived calcium transients in nascent oligodendrocyte processes, potentially revealing a link between localized calcium signaling and actin polymerization. Lastly, adolescent mice (*Mcoln1^-/-^*), in which TRPML1 had been deleted, had significantly impaired myelination and decreased numbers of mature oligodendrocyte, which was associated with a reduction in staining for the phosphorylated form of the actin regulator, PAK1, in the motor cortex and corpus callosum as evidence of decreased TRPML1/Rac1/PAK1 signaling. Together, our work reveals lysosomal TRPML1 activity as a central regulator of oligodendrocyte morphology independent of myelin protein expression and may provide mechanistic insight into the distinct but coordinated pathways that lead to oligodendrocyte differentiation and how lysosomal dysfunction impacts these processes in diseased states.

## Introduction

Oligodendrocyte precursor cells (OPCs) and oligodendrocytes must undergo dramatic cytoskeleton rearrangement to differentiate and produce myelin^1,2^. Differentiating oligodendrocytes have periodic cytoskeletal patterns of actin and spectrin, an actin binding protein, suggesting that actin cytoskeleton in these cells is tightly regulated^3^. The actin cytoskeleton also mediates numerous changes in OPCs and oligodendrocytes, including OPC migration^4^, surveillance of local environments^5^, and active myelin wrapping^6,7^. These dynamic actin cytoskeletal changes are defined by a precisely controlled interplay between the assembly of filamentous actin (F-actin), through polymerization of globular actin (G-actin) subunits, and their disassembly, mediated by F-actin severing and depolymerizing factors^8^. Evidence, thus far, implicates that F-actin polymerization and stabilization are key drivers for OPC migration^9^, establishment of a complex process network^10,11^, and initiation of myelination^6,7^; on the other hand, myelin wrapping around the axon is driven by predominantly F-actin depolymerization and disassembly^7^. Alterations in F-actin are highly sensitive to changes in intracellular calcium (Ca^2+^)^12,13^ and a recent study demonstrated that oligodendrocytes utilize calcium-related cytoskeletal assembly to shape nascent myelin sheath morphology^14^. While there are numerous calcium-permeable receptors and channels on the oligodendrocyte plasma membrane that aid in the assembly of myelin sheaths, lysosomes, which also contain intracellular calcium stores, have not been linked to actin cytoskeletal dynamics in OLs.

Lysosomes are membrane-bound vesicles that have long been known to play a key role in the degradation and recycling of nutrients via endocytosis, phagocytosis, and autophagy^15^. It has become increasingly clear that lysosomes have other important roles, including insertion of the major myelin protein, proteolipid protein (PLP) into the growing myelin membrane^16–18^, and Ca^2+^ signaling^19^. The lysosome is the second largest Ca^2+^ store within the cell, with free [Ca^2+^] of ∼500-600 µM, similar to the endoplasmic reticulum^20^. While there is still debate as to how Ca^2+^ enters the lysosome, there have been a plethora of ion channels identified as functioning to release Ca^2+^, including two pore channel 2 (TPC2) and transient potential receptor mucolipin 1 (TRPML1)^19^. TRPML1, which is localized to late endosomes and lysosomes, is the most well-characterized of these channels, where it plays essential roles in lysosomal exocytosis^21–23^, biogenesis^24^, and positioning^25^. Additionally, TRPML1 has been shown to control the organization of actin cytoskeleton during dendritic cell migration^26^ though this has not been explored within the oligodendrocyte lineage.

Highlighting the potential importance of this channel in oligodendrocyte maturation, inactivating mutations of TRPML1 result in the rare autosomal recessive disease known as type IV mucolipidosis (MLIV)^27,28^. Despite ubiquitous expression of TRPML1 across all organ systems, MLIV primarily affects the central nervous system (CNS), where it is characterized as a hypomyelinating leukodystrophy that manifests as severe psychomotor developmental delay^29^. Data from human imaging^30^ and transgenic mice demonstrate significant myelin involvement and reduction in mature oligodendrocyte number in MLIV^31^, as well as a decreased expression of several mature oligodendrocyte mRNAs, including myelin-associated glycoprotein precursor (*Mag*), myelin basic protein (*Mbp*), myelin-associated oligodendrocyte basic protein (*Mobp*), and proteolipid protein (*Plp*)^32^. Despite these data, there has been minimal investigation into the role of TRPML1 in regulating oligodendrocyte maturation and myelination during development.

The spontaneous assembly of F-actin is an inherently inefficient process; however, to overcome this, cells utilize membrane-bound factors that can directly nucleate actin, including actin-related protein 2/3 (Arp2/3) complex^33^. While actin nucleation and filament generation have been associated with activity at the plasma membrane to facilitate cell migration, endocytic structures, and lamellipodia/filopodia formation^33^, there is accumulating evidence that actin dynamics also occur on the surface of intracellular organelles, including mitochondria and lysosomes^34,35^. Nucleation factors, like Arp2/3, are recruited to membranes via several upstream signaling pathways, including the small GTPase Ras-related C3 botulinum toxin substrate 1 (Rac1). In its active GTP-bound form Rac1 influences the cytoskeleton through two distinct pathways. The first modulates actin nucleation and branching through the WAVE regulatory complex and Arp2/3 activation, while the second inhibits the actin-disassembly factor Cofilin-1 through activation of PAK1 and LIMK1^36^. Both Arp2/3 and PAK1 have been shown to positively regulate oligodendrocyte morphology through changes in the actin cytoskeleton^7,37^; however, the upstream events that drive these signaling cascades have not been well-defined with differentiating OLs.

Here, for the first time, we report that as oligodendrocytes mature, endolysosomes containing TRPML1 migrate into nascent processes where they are poised to influence intracellular signaling. In line with this, we demonstrate that TRPML1 controls actin polymerization through the Rac1/PAK pathway, a known regulator of the cytoskeleton. Additionally, spontaneous lysosome-derived calcium transients occur within the oligodendrocyte lineage, and these happen almost exclusively within developing processes. Knockout of TRPML1 in adolescent mice (*Mcoln1^-/-^)* results in impaired developmental myelination and reduced phosphorylation of PAK1 within oligodendroglia lineage cells suggesting that actin assembly is not being appropriately regulated in the absence of TRPML1. Together, these data highlight the essential role that lysosomes play in morphologic changes in oligodendrocytes during development via TRPML1.

## Results

### Endolysosomes localize into developing oligodendrocyte processes

TRPML1 is exclusively expressed on endolysosomal membranes^38^ across all cell types; thus, we sought to determine where endolysosomes are located in cells at different stages of maturation in the oligodendrocyte lineage. Primary rat OPC cultures were either maintained in growth medium or switched to differentiation medium (DM) for 24, 48, or 72 hours prior to being incubated with the cell-permeable dye, LysoTracker Green, which accumulates in acidic compartments, including late endosomes and lysosomes, allowing real-time, live tracking. During the OPC stage, LysoTracker Green positive puncta (LysoTracker Green^+^) were found primarily in the perinuclear region with few puncta trafficking out into nascent processes over a 10-minute live imaging period (Fig. 1a and Supplementary Video 1). As OPCs differentiated over a 72-hour period, a portion of LysoTracker Green^+^ endolysosomes migrated into oligodendrocyte processes with a substantial proportion remaining in the soma (Fig. 1a and Supplementary Video 2). To quantify the impact of maturation stage on endolysosome motility, we tracked individual endolysosomes within a 50 μm segment and measured mean speed and cumulative distance traveled. Endolysosomal mean speed and cumulative distance traveled increased through 48 hours of differentiation and returned to OPC levels by 72 hours (Fig. 1b-c). Interestingly, while endolysosomal speed and distanced traveled were greatly reduced at three days of maturation, endolysosomes retained localization within oligodendrocyte processes, raising the possibility that they may serve as signaling hubs, such as those required for actin nucleation and disassembly.

**Figure 1.**
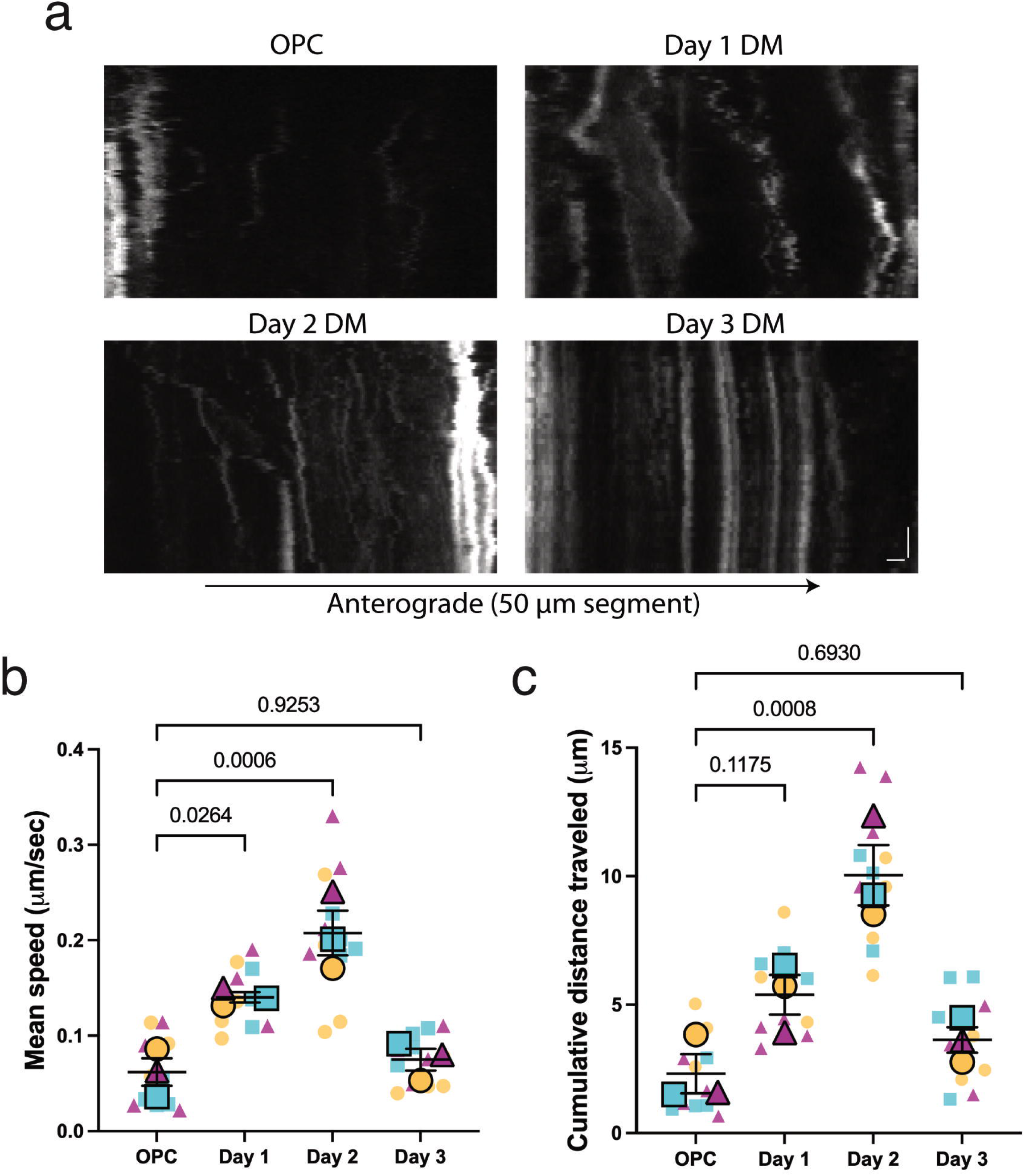
Endolysosomes localize into developing oligodendrocyte processes. **(a)** Representative kymographs of LysoTracker+ puncta in primary processes of OPCs and oligodendrocytes at different stages of differentiation. Vertical scale bar represents 1 minute and horizontal scale bar represents 5 μm. **(b)** Endolysosome mean speed significantly increased over two days of differentiation and returned to OPC levels by three days of maturation. **(c)** Similar to mean speed, endolysosome cumulative distance significantly increased through two days of differentiation and returned to baseline OPC levels at day three of differentiation. N=3 biological replicates (preps); small symbols represent individual cells; each shape and color are from a different biological replicate. One-way ANOVA followed by Tukey’s post-hoc test.

Since TRPML1 is implicated in endolysosomal trafficking via the efflux of Ca^2+^, recruitment of the Ca^2+^ sensor ALG2, and association with the motor protein dynein^25^, we examined whether oligodendrocyte endolysosomal motility was affected by activation or inhibition of TRPML1. OPCs that underwent differentiation for 24 to 48 hours were labeled with LysoTracker Green and treated with either the TRPML1 agonist, MLSA1, or the TRPML1 antagonist, MLSI1, for 10 minutes prior to imaging. As expected, activation of TRPML1 resulted in increased endolysosomal motility (mean speed and cumulative distance traveled), while inhibition resulted in stalling of endolysosomes (Supplementary Fig. 1a-d). Additionally, when we quantified the percentage of time each endolysosome moved (retrograde or anterograde) or remained stationary, we saw a greater proportion of MLSI-treated endolysosomes remaining stationary (Supplementary Fig. 1e-f). Taken together, these data demonstrate that during oligodendrocyte differentiation endolysosomes migrate into nascent processes and remain there once cells have matured. Importantly, we demonstrated that TRPML1 is expressed throughout the oligodendrocyte lineage and activity of this channel impacts endolysosome motility in developing OLs.

### Spontaneous calcium events via TRPML1 occur almost exclusively in OPC and oligodendrocyte processes

TRPML1 has been characterized as a non-selective cation channel, permeable to Ca^2+^, Fe^2+^, Zn^2+^, Na^+^, and K^+^; however, it primarily functions as a Ca^2+^ efflux channel. To measure endogenous TRPML1 activity and Ca^2+^ efflux through this channel, we engineered GCaMP6s, a single-wavelength calcium indicator, to the cytoplasmic amino terminus of TRPML1 under an *Olig2* promoter to ensure expression within the oligodendrocyte lineage. When transfected into OPCs, GCaMP6s-TRPML1 was mainly localized to LysoTracker-Red positive compartments (Supplementary Fig. 2).

Intriguingly, under basal conditions, we observed spontaneous lysosome-derived Ca^2+^ events in the absence of MLSA1 treatment (Fig. 2a and Supplementary Video 3), as well as large calcium events upon addition of MLSA1 (Supplementary Video 4). These calcium events could be classified into two distinct categories. Transients that were identified as “fast” appeared and dissipated within an approximately three second period and were restricted to a localized zone within the process. “Prolonged” transients, on the other hand, occurred over an approximately nine to ten second period and typically filled the entire process (Fig. 2a-b). Importantly, these calcium events, whether they were fast or prolonged, occurred almost exclusively in the processes rather than the soma (Fig. 2c). These data demonstrate that endogenous TRPML1 activity occurs within the OL lineage and calcium efflux from TRPML1 appears to be restricted to their processes.

**Figure 2.**
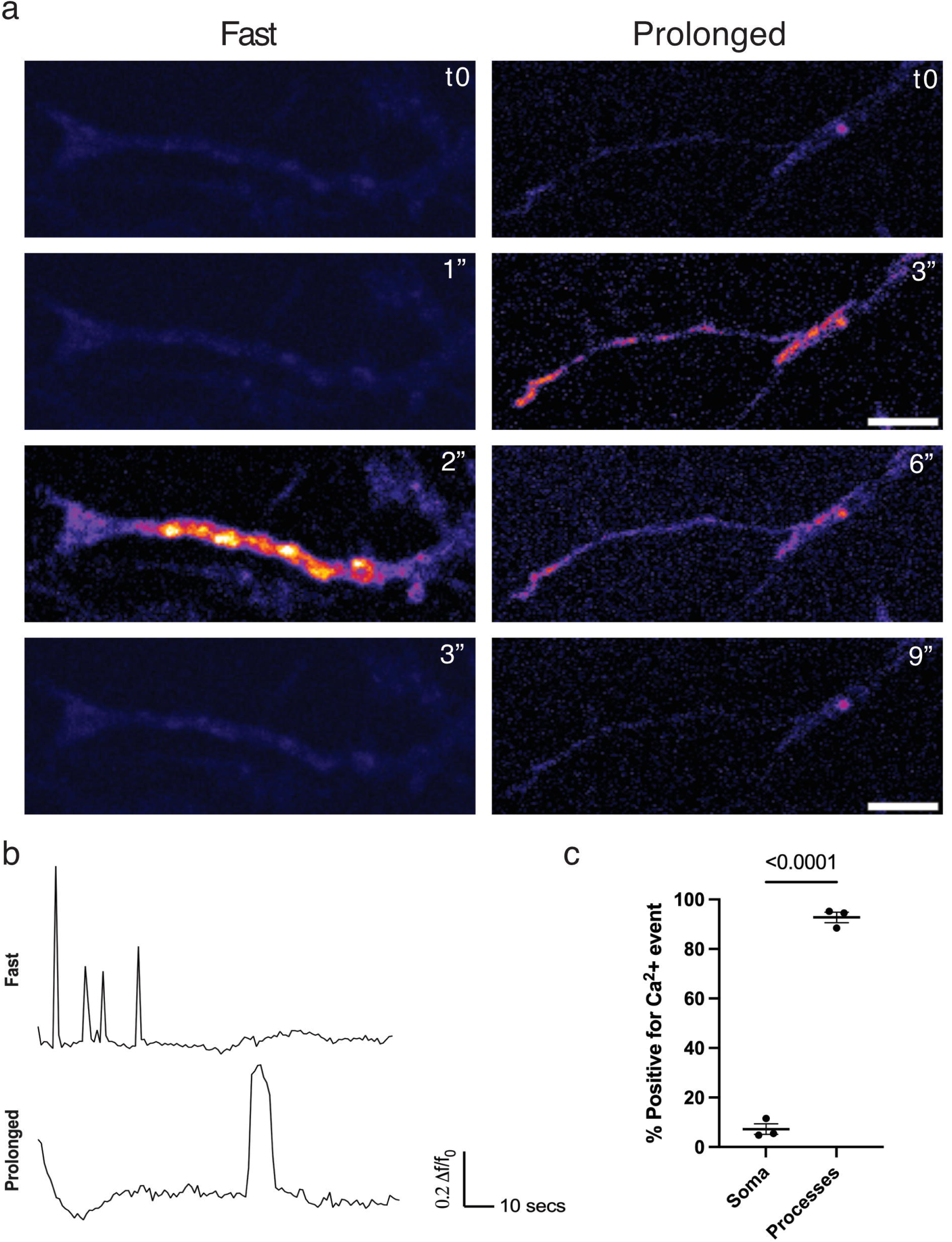
Spontaneous calcium efflux through TRPML1 predominantly occurs in OPC and oligodendrocyte processes. **(a)** Representative images of two different calcium dynamics observed in the processes of OPCs. The first (labeled as “fast”) appears and dissipates over a three second period, with the calcium burst lasting for one second. The second (labeled as “prolonged”) one occurs over a six second period and proceeds along the entire length of the process. Scale bar = 10 µm. **(b)** Example traces of fast and prolonged calcium events shown in (a). **(c)** Quantification of calcium (≥ 0.2 Δf/f_0_) events occurring in the soma and processes of oligodendrocyte lineage cells. N=3 biological replicates. Unpaired two-tailed Student’s *t* test.

### TRPML1 controls oligodendrocyte morphology during differentiation

Individuals with mutations in *Mcoln1* display severe hypomyelination, suggesting a deficit in oligodendrocyte maturation^27^. To determine whether manipulation of TRPML1 activity during oligodendrocyte differentiation impacted the ability of OPCs to become mature oligodendrocytes, we treated primary rat OPCs with either MLSA1 or MLSI1 at the time of differentiation induction and assessed stage-specific lineage markers: A2B5 (OPCs), galatocerebroside (GalC; immature/mature oligodendrocytes), and proteolipid protein (PLP; mature oligodendrocytes) (Fig. 3a). We did not observe any significant differences in the percentage of cells expressing oligodendrocyte lineage markers following activation or inhibition of TRPML1 during oligodendrocyte maturation (Fig. 3b-d) and quantification of DAPI+ nuclei indicated no change in cell viability across the treatments (Fig. 3e). Further, overexpression or knockdown of TRPML1 by lentiviral infection also had no impact on lineage marker expression after three days of differentiation (Supplementary Fig. 3). However, while there were no changes in oligodendrocyte lineage marker expression, there were striking morphologic alterations observed (Fig. 3a). To further investigate this, we reconstructed individual PLP^+^ mature oligodendrocytes using the SNT plugin in ImageJ and analyzed these cells for morphologic complexity and primary process length (Fig. 4a). Cells treated with MLSI1 during differentiation had a significant reduction in process branching compared to vehicle-treated cultures, especially at distances more distal to the soma (> 35 μm; Fig. 4b-c). Conversely, activation of TRPML1 resulted in greater morphologic complexity in mature, PLP+ oligodendrocytes (Fig. 4b-c). Additionally, we also measured the length of primary processes originating from the soma of PLP+ mature oligodendrocytes. In line with the reduction in branching complexity, inhibition of TRPML1 induced shorter primary process length (Fig. 4d). Taken together, these data demonstrate that TRPML1 plays an essential role in regulating oligodendrocyte morphology during the early stages of differentiation.

**Figure 3.**
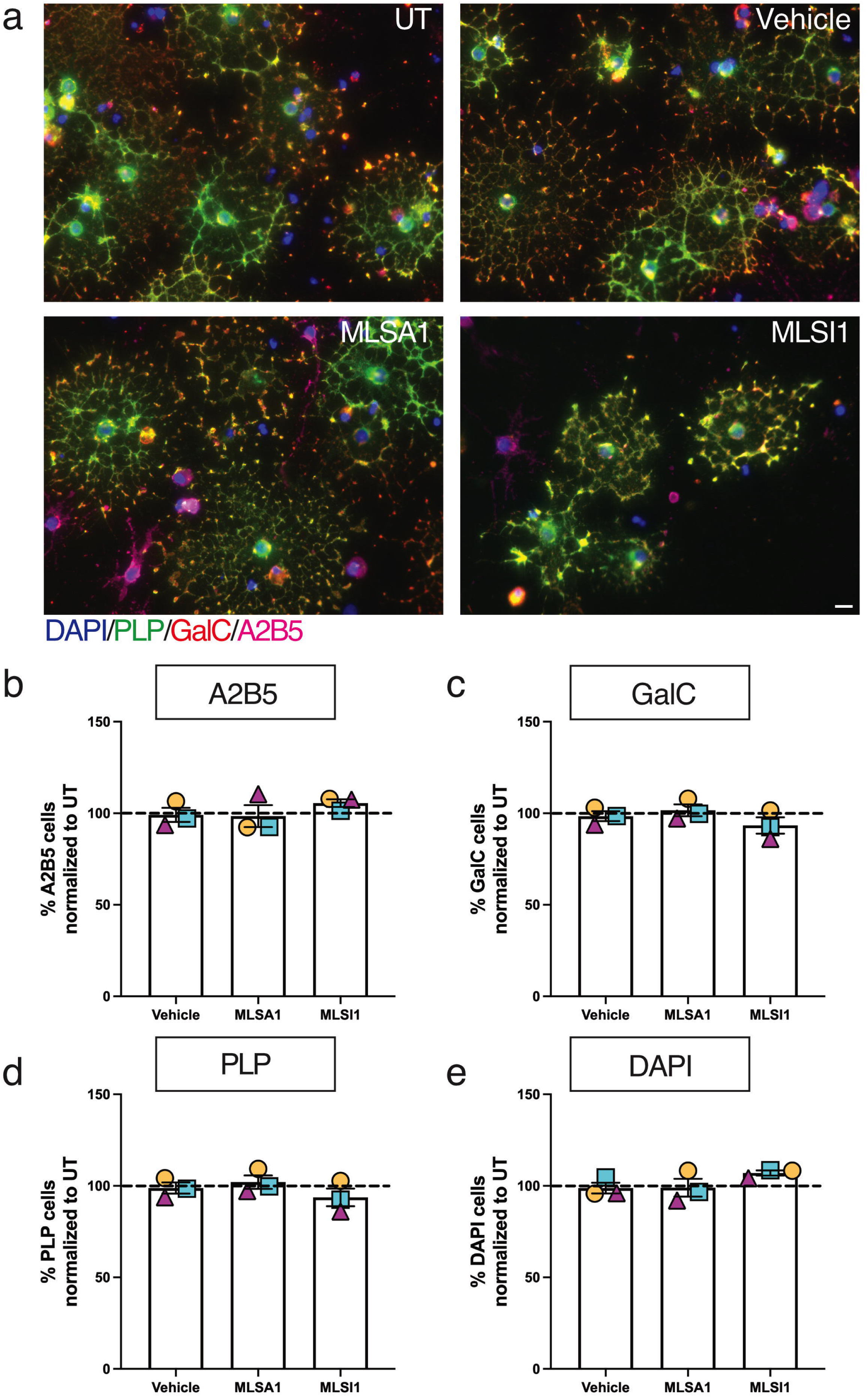
Activation or inhibition of TRPML1 during differentiation does not alter expression of oligodendrocyte lineage markers. **(a)** Representative images of three-day differentiated oligodendrocyte cultures treated at the time of differentiation with either vehicle (DMSO), MLSA1 (TRPML1 agonist; 20 μM), or MLSI1 (TRPML1 antagonist; 15 μM) and stained for A2B5 (OPCs), GalC (immature/mature oligodendrocytes), PLP (mature oligodendrocytes), and DAPI (nuclei). Scale bar = 50 μm. **(b)** No changes were observed in the percentage of cells expressing the OPC marker, A2B5, across all treatment groups. **(c)** There were no significant alterations in the percentage of cells expressing the immature/mature oligodendrocyte marker, GalC, across all treatment groups. **(d)** As observed with A2B5 and GalC, there were no significant differences in the percentage of cells labeled with the mature oligodendrocyte marker, PLP, across all groups. **(e)** Total cell number, as measured by DAPI+ nuclei staining, was not significantly different across any of the treatment groups. N=3 biological replicates (preps). One-way ANOVA followed by Tukey’s post-hoc test; all p>0.05.

**Figure 4.**
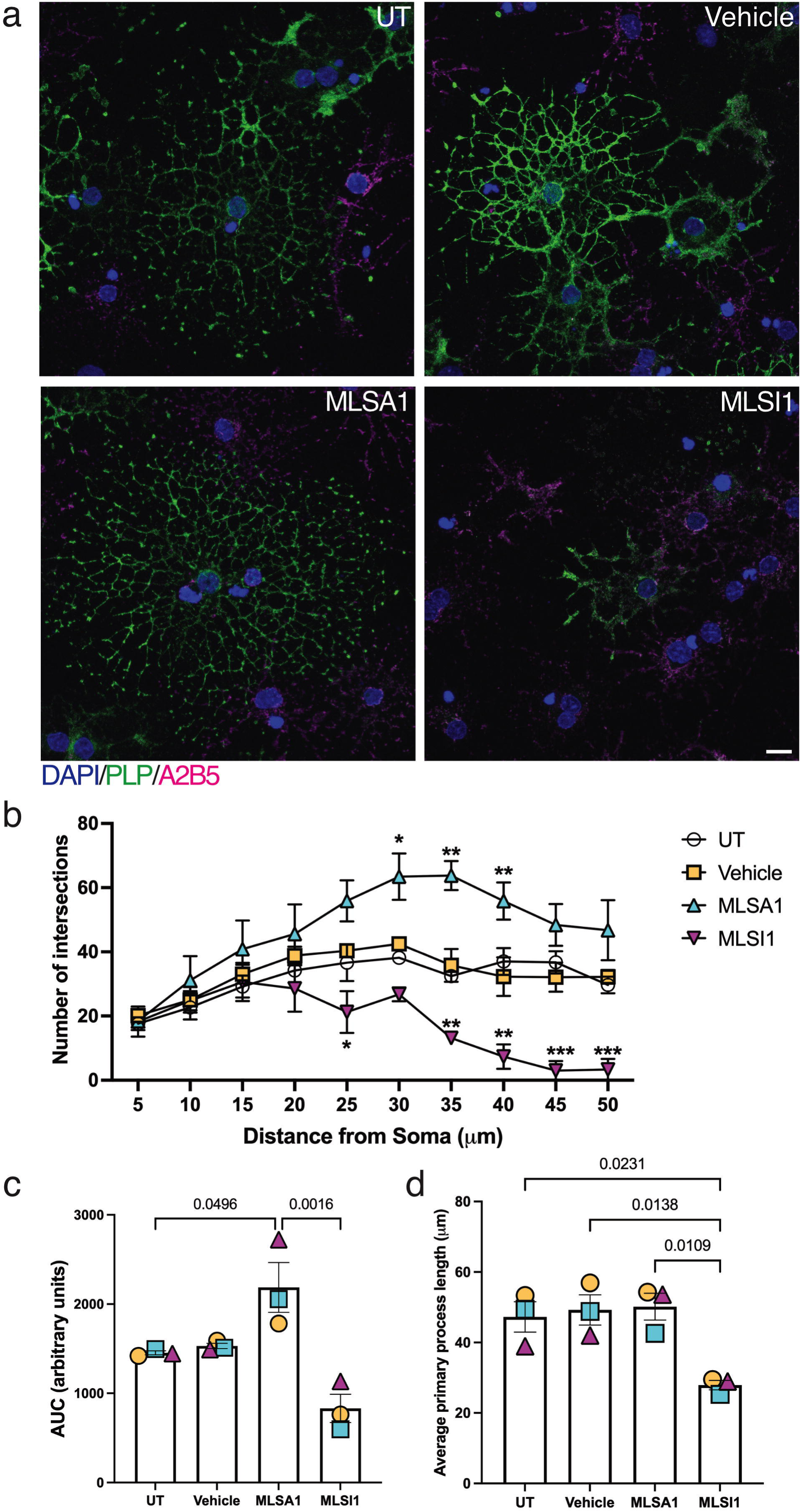
TRPML1 controls oligodendrocyte morphology during differentiation. **(a)** Representative images of three-day differentiated oligodendrocyte cultures treated at the time of differentiation with either vehicle (DMSO), MLSA1 (20 μM), or MLSI1 (15 μM); scale bar = 10 μm. **(b)** MLSA1-treated PLP+ cells exhibited significantly greater morphologic complexity as measured via Sholl analysis. Conversely, MLSI1-exposed cells had simpler morphology, particularly at distal regions of the processes. **(c)** Quantification of Sholl analysis by area under the curve. **(d)** Primary process length (processes originating from the soma) was significantly decreased in MLSI1-treated cultures. N=3 biological replicates (preps). Two-way ANOVA or one-way ANOVA followed by Tukey’s post-hoc test; *p<0.05, **p<0.01, ***p<0.001.

To determine whether TRPML1 also modulates oligodendrocyte lineage marker expression and morphology during myelination initiation, we used an artificial nanofiber system to model *in vitro* myelination^39^. Interestingly, in comparison to 2D monolayer culture, inhibition of TRPML1 during myelination initiation significantly increased the percentage of A2B5+ cells, while reducing the number of PLP+ cells (Supplementary Fig. 4a-d). Additionally, as observed in our differentiation paradigm, inhibition of TRPML1 had a significant impact on oligodendrocyte morphology, with reductions in branching complexity, average primary process length, and number of myelinating processes (Supplementary Fig. 5a-d). Cells treated with MLSA1 had subtle, but significant, alterations in morphologic complexity, while those treated with MLSI1 had overt decreases in branching with no processes found at 45 or 50 μm away from the soma (Supplementary Fig. 5b). This was also reflected in the average primary process length, with agonist-exposed cultures exhibiting significantly longer primary processes and inhibitor-treated ones with shorter ones (Supplementary Fig. 5c). Lastly, we analyzed the number of processes interacting with artificial nanofibers (“myelinating processes”) and found that nanofiber cultures treated with MLSI1 had significantly fewer myelinating processes compared to all other groups (Supplementary Fig. 5d), suggesting TRPML1 regulates myelinating process length, number, and complexity, a function not previously described.

### TRPML1 alters actin filament levels during differentiation

Given the relationship we observed between process extension and TRPML1 and reports from other groups demonstrating that the actin cytoskeleton controls oligodendrocyte myelination via two distinct steps, the first of which drives oligodendrocyte process outgrowth and branching via actin assembly^7,14^, we sought to determine the impact of TRPML1 on actin filament levels and polymerization during oligodendrocyte maturation. OPC cultures were treated at the time of differentiation with either MLSA1 or MLSI1 for 24, 48, or 72 hours and stained with phalloidin, a peptide that binds with high affinity and specificity to filamentous actin (F-actin) (Fig. 5a). We observed a significant increase in actin filament levels after 24 hours of MLSA1 treatment and this decreased over 48 and 72 hours, with the 72-hour timepoint significantly less than untreated and vehicle (Fig. 5b-d). Conversely, cultures treated with MLSI1 had a robust reduction in actin filament levels across all three timepoints analyzed (Fig. 5b-d), reflecting the significant morphologic disruption we observed (Fig. 3 & 4). We next complemented our pharmacologic studies with genetic knockdown or overexpression of TRPML1 via lentiviral infection (Supplementary Fig. 6a-b). As observed with activation of TRPML1, lentiviral-mediated overexpression of TRPML1 increased F-actin content 24 hours after induction of differentiation; conversely, knockdown of TRPML1 via *Mcoln1* shRNA significantly inhibited levels of F-actin within differentiating oligodendrocytes (Supplementary Fig. 6a-b).

**Figure 5.**
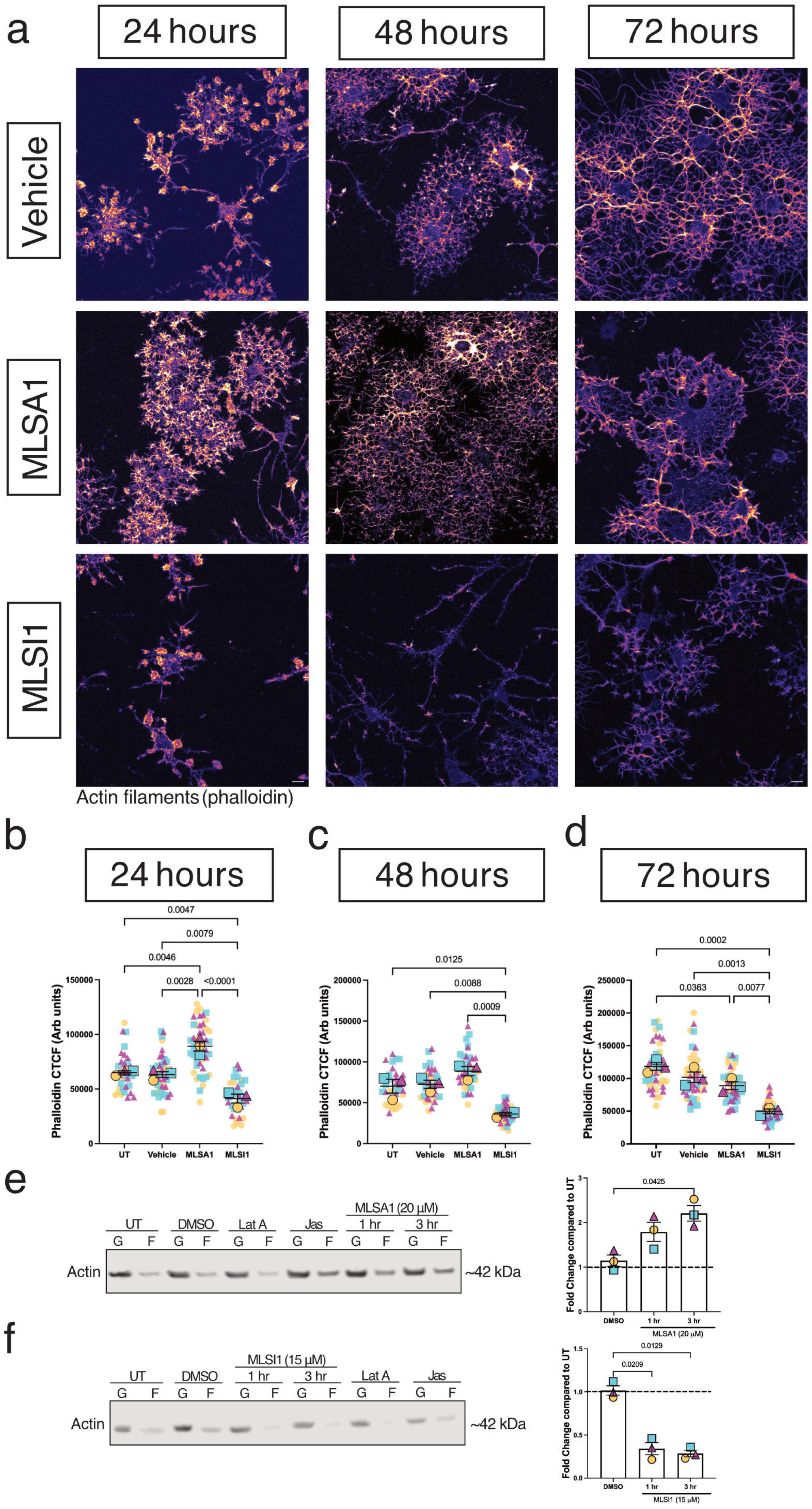
TRPML1 alters actin filament levels during oligodendrocyte maturation. **(a)** Representative images of vehicle-, MLSA1-, or MLSI1-treated cultures at one, two, or three days after differentiation and stained with phalloidin; scale bar = 10 μm. **(b)** 24-hour treatment with MLSA1 induced a significant increase in F-actin content in oligodendrocyte cultures, while MLSI1-treated cultures had a significant reduction in F-actin filament levels. **(c)** MLSI1-treated oligodendrocyte cultures had significantly less F-actin content after 48 hours of treatment compared to all other treatment groups. **(d)** 72 hours after treatment, both MLSA1- and MLSI1-treated cultures had a reduction in F-actin filament levels compared to untreated cultures. **(e)** Three-hour treatment with MLSA1 increased the ratio of F-actin to G-actin, indicating a shift towards actin polymerization. **(f)** Treatment with MLSI1 for one or three hours significantly decreased the F/G-actin ratio, indicating a shift towards actin depolymerization. N=3 biological replicates (preps); small symbols represent individual cells; each shape and color are from a different biological replicate. One-way ANOVA followed by Tukey’s post-hoc test.

Based on the early changes in actin filament levels as seen with phalloidin staining, we would expect a shift in the ratio of F-actin to globular actin (G-actin) in favor of F-actin in response to activation of TRPML1. Differentiating OPC cultures treated at the time of differentiation with MLSA1 for three hours exhibited a 2.5-fold increase in the F/G actin ratio relative to vehicle-treated controls, indicative of a shift towards increased actin polymerization (Fig. 5e). This functional change in the actin ratio is consistent with the time-dependent increase in actin filament staining induced by MLSA1 (Fig. 5a). Conversely, treatment of maturing OPC cultures with MLSI1 for the same timepoints led to a significant reduction in the F/G actin ratio, suggesting an increase in actin disassembly (Fig. 5f). As before, these changes in the F/G actin ratio are in line with the time frame observed for actin filament levels where changes are observed with MLSI1 as early as six hours post treatment (data not shown).

The actin cytoskeleton is a major cellular target of calcium and TRPML1, while permeable to a number of cations, is regarded primarily as a lysosomal calcium efflux channel^40,41^. Thus, we sought to clarify whether the effects on the actin cytoskeleton were specific to TRPML1 or could potentially be induced by calcium efflux through other lysosomal channels. We chose to investigate two-pore channel 2 (TPC2), as it is expressed within OPCs and calcium release through this channel has been well documented^42–44^. Utilizing a cell-permeable TPC2 agonist that mimics NAADP action, TPC2-A1-N^45^, as well as a TPC2 antagonist, SG-094^46^, we stained differentiating OPC cultures following 24 or 48 hours of treatment with phalloidin (Supplementary Fig. 7a). As previously observed, activating TRPML1 led to a robust increase in actin filament levels while inhibition of TRPML1 suppressed actin filament formation at both time points (Supplementary Fig. 7b-c). Surprisingly, treatment with either TPC2-A1-N or SG-094 had no discernable effect on actin filaments during the early stages of differentiation (Supplementary Fig. 7b-c), suggesting that lysosomal calcium release alone is not sufficient to induce alterations in actin filament content and that TRPML1 controls actin polymerization through either a calcium-independent mechanism or via specific signaling partners the channel is coupled to on the lysosomal membrane.

### TRPML1 activates the Rac1/PAK pathway in the oligodendrocyte lineage

To investigate the link between TRPML1 activation and actin polymerization, we examined the role of the small GTPase Rac1, a well-described master regulator of actin polymerization previously identified as a TRPML1 binding partner^47^. We first sought to determine whether, based on the proposed interaction between TRPML1 and Rac1 on the endolysosomal surface, activation of TRPML1 would modulate levels of Rac1 bound to GTP (Rac1-GTP). Differentiating OPC cultures were treated with MLSA1, and lysates were subjected to pulldown with PAK1 agarose beads. We observed a significant increase in Rac1-GTP levels following 15- or 30-minutes of TRPML1 treatment; furthermore, we did not see any changes into total Rac1 levels, suggesting that TRPML1 activity specifically regulates the amount of Rac1 bound to GTP (Fig. 6a).

**Figure 6.**
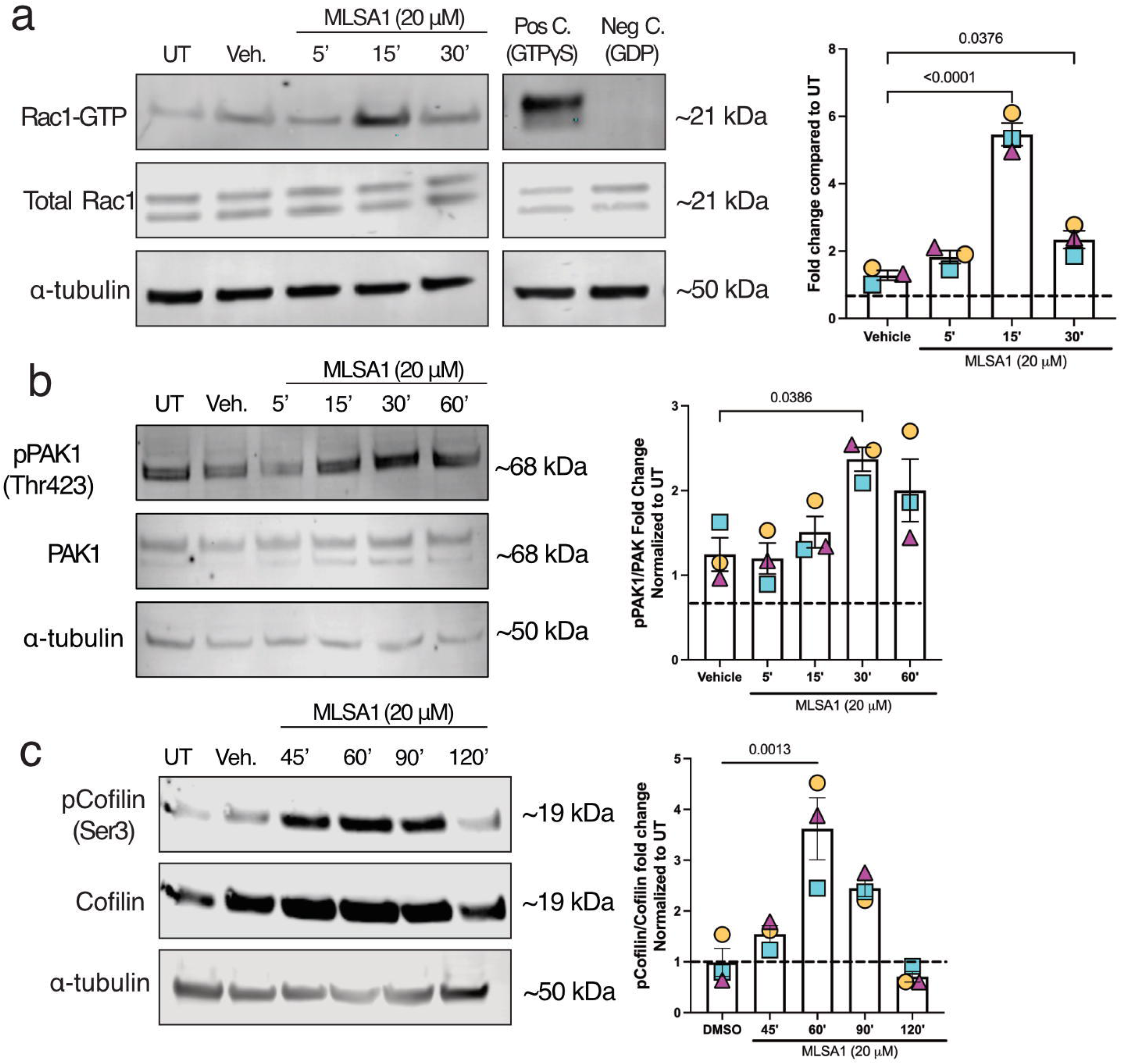
TRPML1 activates the Rac1/PAK pathway in oligodendrocytes. **(a)** Differentiating oligodendrocyte cultures treated with MLSA1 exhibited a significant increase in Rac1-GTP after 15 and 30 minutes of treatment. **(b)** Increased levels of phosphorylated PAK1 were observed after 30 minutes of MLSA1 treatment. **(c)** Levels of phosphorylated cofilin were elevated following one hour of MLSA1 treatment in differentiating oligodendrocyte cultures. N=3 biological replicates (preps). One-way ANOVA followed by Tukey’s post-hoc test.

Next, we investigated phosphorylation of proteins downstream of Rac1 that influence actin cytoskeletal dynamics in TRPML1 treated differentiating OPCs. We observed significant increases in phosphorylation of PAK1 and Cofilin-1 in a time-dependent manner, with peak pPAK1 at 15 minutes and peak Cofilin-1 at 1 hour (Fig. 6b-c), in line with previously published reports regarding the time course of phosphorylation events on these proteins^48^. This signaling pathway culminates with the phosphorylation and inactivation of Cofilin, which acts as an actin-severing protein and promotes the dissociation of F-actin to G-actin. These findings suggest TRPML1 activation results in Rac1/PAK pathway activation in oligodendroglia which has implications for how essential actin-based morphologic changes occur during oligodendrocyte maturation.

Since we have identified Rac1 as the protein bridging TRPML1 to alterations in the cytoskeleton, we next sought to determine if Rac1 activity was necessary for the increase in F-actin content within oligodendrocytes. First, we determined that pretreatment of OPC cultures with the specific Rac1 inhibitor, NSC23766^49^ (100 ug/mL; 15 minutes) did not alter the percentage of cells expressing oligodendrocyte lineage markers after three days of differentiation, in the presence or absence of MLSA1 (Supplementary Fig. 8a-e). However, pretreatment of OPC cultures with NSC23766 prior to the addition of MLSA1 completely blocked the ability of the TRPML1 agonist to increase the amount of F-actin at the early stages of differentiation (Fig. 7a-c). Together, these data suggest activation of TRPML1 changes the actin cytoskeleton through modulation of the Rac1/PAK pathway.

**Figure 7.**
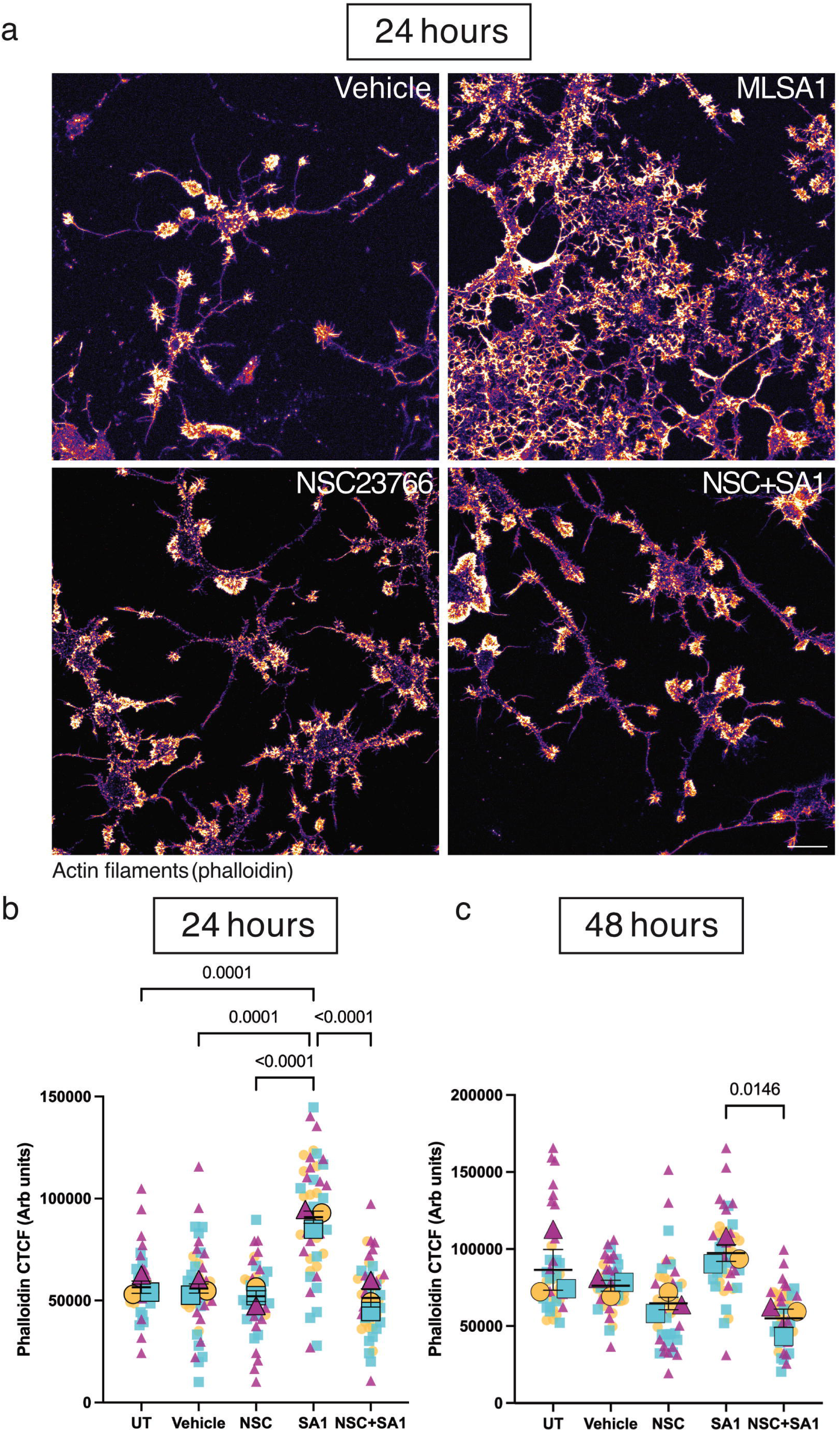
Blockade of Rac1 activation prevents TRPML1-mediated effects on F-actin content in differentiating oligodendrocytes. **(a)** Representative images of one-day differentiated oligodendrocytes treated with vehicle-only, MLSA1-only, NSC23766-only, or NSC23766 with MLSA1 (abbreviated NSC + SA1), and stained with phalloidin to visualize actin filaments; scale bar = 10 μm. **(b)** Pre-treatment of oligodendrocytes with NSC23766 prior to MLSA1 application completely prevented the alterations in F-actin content induced by MLSA1 alone after one day of differentiation. No significant changes were observed with NSC23766 alone. **(c)** F-actin content was significantly reduced NSC23766+MLSA1 treated cultures compared to MLSA1-only in oligodendrocytes differentiated for two days. N=3 biological replicates (preps); small symbols represent individual cells; each shape and color are from a different biological replicate. One-way ANOVA followed by Tukey’s post-hoc test.

### *Mcoln1^-/-^* mice have impaired myelination and phosphorylation of PAK1 within oligodendroglia

Patients with mutations in *MCOLN1*, the gene that encodes for TRPML1, develop a rare autosomal recessive lysosomal storage disorder known as mucolipidosis type IV (MLIV). Previous data from *Mcoln1^-/-^*mice revealed that mutant mice had a significantly thinner corpus callosum at two and seven months of age and this coincided with decreased PLP staining intensity^31,50^, suggestive of a developmental phenotype with impaired oligodendrocyte maturation. We sought to confirm this in 21-day old pups, a time point associated with the onset of peak oligodendrocyte maturation and myelination in the developing mouse brain^51^. First, myelination was assessed in the motor cortex and medial corpus callosum by staining for PLP (Fig. 8a,d). Mutant mice (*Mcoln1^-/-^*) exhibited significantly decreased PLP staining intensity within the medial corpus callosum compared to both wild-type (*Mcoln1^+/+^*) and heterozygous (*Mcoln1^+/^*) mice (Fig. 8b). Interestingly, PLP staining intensity was slightly, but significantly, higher in heterozygous mice compared to wild-type. Within the motor cortex, similar effects were observed regarding PLP staining intensity. Since myelination of the deeper layers (IV-V) of the cortex occur prior to the superficial ones (I-III), we assessed myelination with PLP staining in both superficial and deep layers of the motor cortex. We observed a significant reduction in PLP fluorescence intensity in both superficial and deep layers of the motor cortex of mutant mice, with more drastic changes observed in the area not as heavily myelinated (layers I-III) (Fig. 8e & Supplementary Fig. 9a). As PLP is expressed by mature, myelinating OLs, we also stained for aspartoacylase (ASPA). Unlike PLP, mutant mice only had subtle changes in ASPA+ cell count in the corpus callosum and layers IV-VI of the motor cortex but not layers I-III of the motor cortex (Fig. 8c,f & Supplementary Fig. 9b).

**Figure 8.**
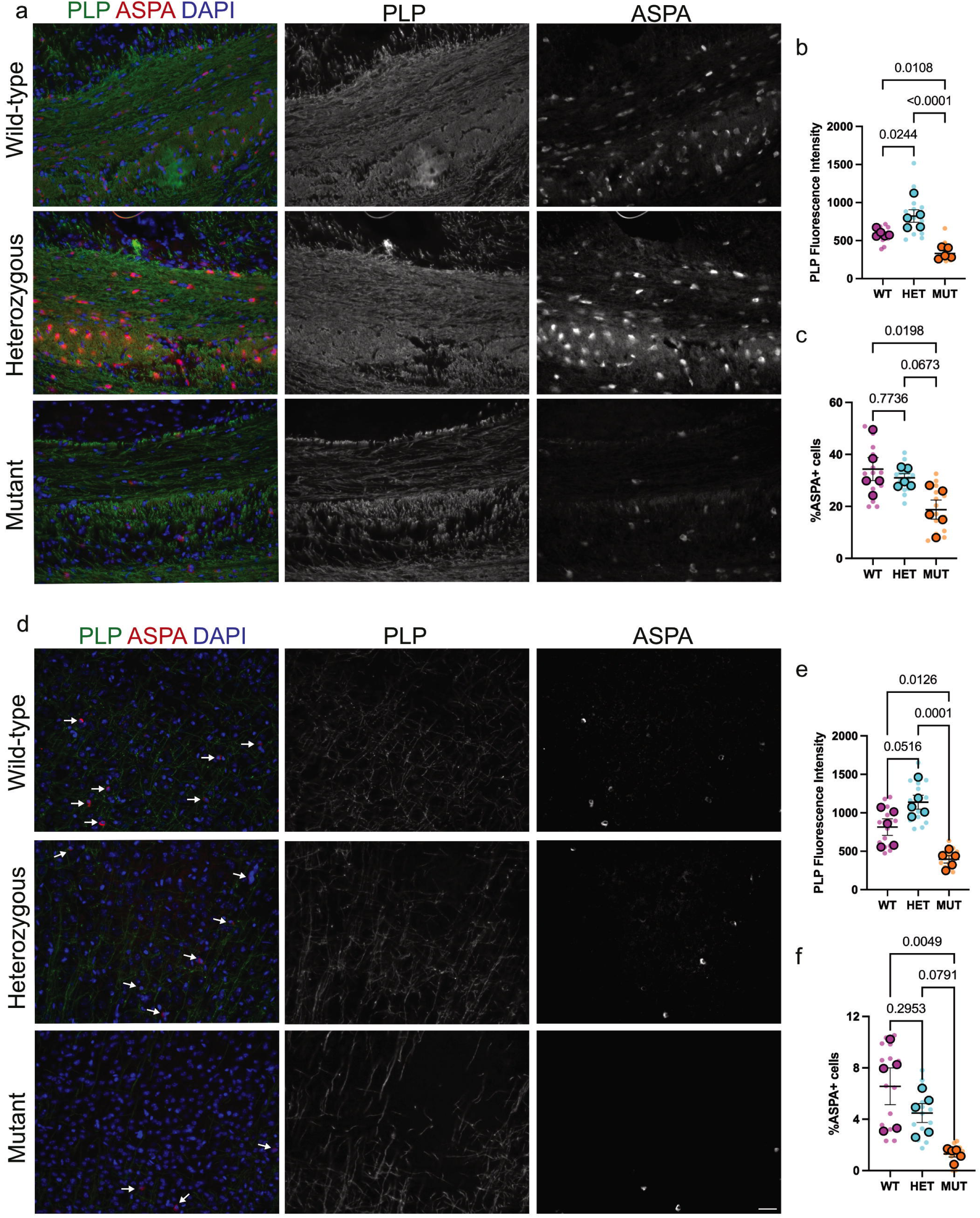
P21 *Mcoln1^-/-^*mice have impaired myelination and fewer mature OLs in the corpus callosum and motor cortex. **(a)** Representative images of the medial corpus callosum stained for ASPA (mature OL), PLP (myelin protein), and DAPI (cell nuclei); scale bar = 50 μm. **(b)** TRPML1 KO mice had a significant reduction in PLP staining intensity compared to wild-type and heterozygous littermates. Heterozygous mice had greater PLP fluorescence signal compared to wild-type mice. **(c)** There were significantly fewer mature OLs in TRPML1 KO when compared to wild-type, but not heterozygous, mice. **(d)** Representative images of the motor cortex (layers IV-VI) stained for ASPA, PLP, and DAPI. **(e)** As observed in the corpus callosum, TRPML1 KO mice had a significant decrease in PLP staining intensity compared to both wild-type and heterozygous mice. **(f)** Similar to the corpus callosum, TRPML1 KO mice had fewer mature OLs compared to wild-type mice, but not heterozygous littermates. N=5 animals per genotype; small symbols represent individual sections. One-way ANOVA followed by Tukey’s post-hoc test.

Our *in vitro* studies elucidated the critical role of Rac1/PAK1 activation in regulating TRPML1-mediated actin polymerization and morphologic changes in oligodendrocytes. Thus, we sought to test whether oligodendroglial lineage cells in TRPML1 knockout mice have reduced phosphorylated PAK1 in the corpus callosum. Immunohistochemical analysis of pPAK1 in Olig2+ cells in the corpus callosum was conducted and multispectral imaging was used to quantify changes in phosphorylation levels in individual oligodendroglial cells. In TRPML1 KO mice, the level of pPAK1 within Olig2+ in the corpus callosum was significantly decreased compared to both wild-type and heterozygous mice, in line with our *in vitro* data demonstrating that TRPML1 is an upstream regulator of Rac1/PAK1 activity in the oligodendrocyte lineage (Fig. 9a-d).

**Figure 9.**
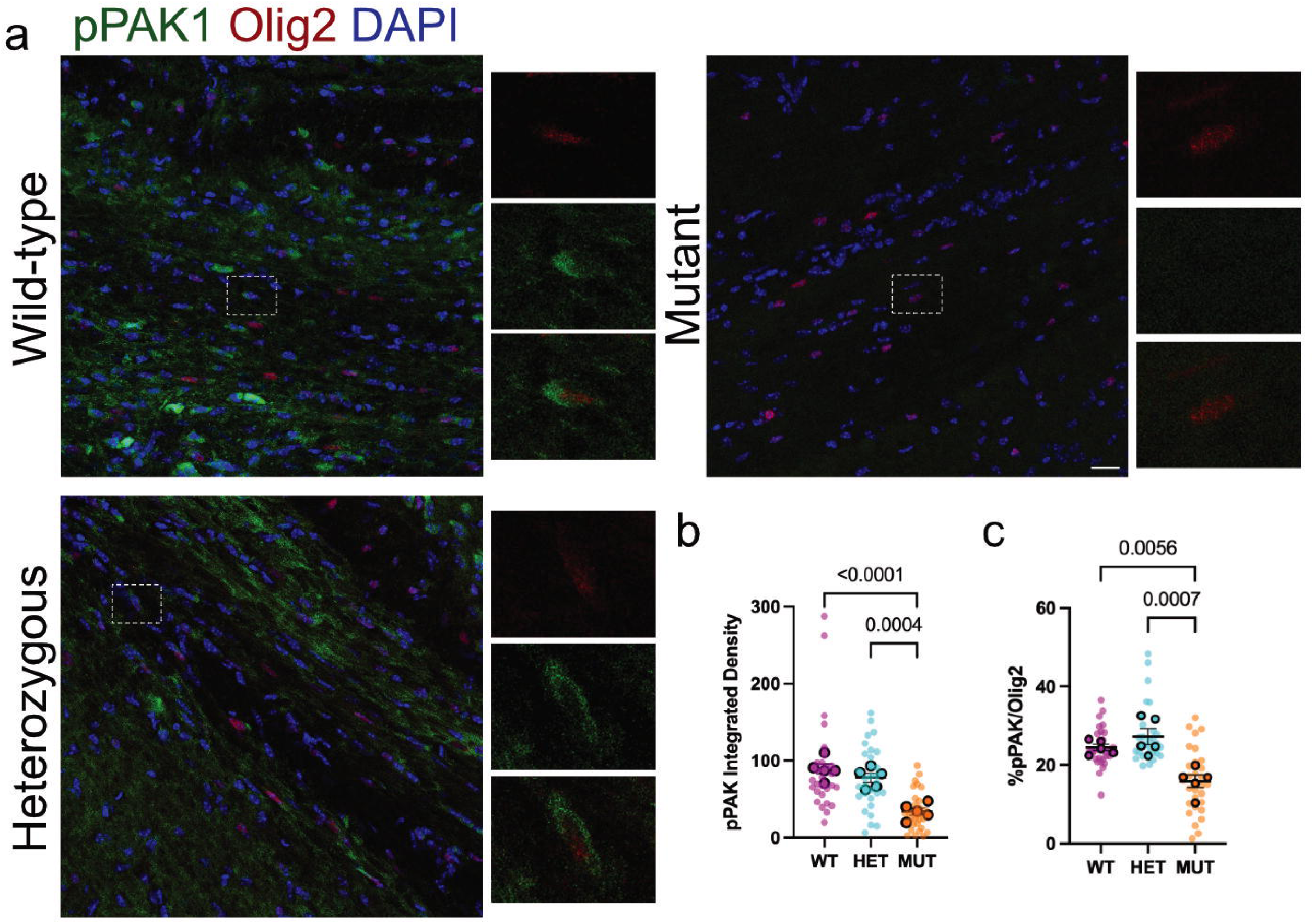
Phosphorylation of PAK1 is disrupted in *Mcoln1^-/-^* mice in oligodendrocyte lineage cells. **(a)** Representative images of the medial corpus callosum stained for pPAK1 (Thr423), Olig2 (OL lineage cells), and DAPI; scale bar = 50 μm. **(b)** Olig2+ cells within the corpus callosum of TRPML1 KO mice had a significant reduction in pPAK1 integrated density compared to wild-type and heterozygous mice. **(c)** The percentage of Olig2+ cells that were also positive for pPAK1 was significantly decreased in TRPML1 KO mice. N=5 animals per genotype; small symbols represent individual sections. One-way ANOVA followed by Tukey’s post-hoc test.

## Discussion

Upon differentiating, OPCs are required to undergo massive alterations to their cytoskeleton to generate processes that eventually engage with and wrap around neuronal axons. Previous evidence has implicated both actin polymerization and calcium dynamics as important regulators of this process; however, the molecular mediators linking these two is unclear. Here, we report a previously unknown mechanism linking lysosome activity to dynamic actin alterations through the cation channel, TRPML1. We found that as OPCs mature into oligodendrocytes, endolysosomes migrate out into nascent processes during the early stages and eventually become stationary. Surprisingly, we found that TRPML1 activity did not alter the ability of oligodendrocytes to express mature myelin proteins, such as MBP or PLP; however, we did observe striking morphologic changes, including process length and branching complexity. These alterations were driven by actin polymerization whereby activation or overexpression of TRPML1 increased F-actin content and inhibition or knockdown resulted in significant decreases in F-actin. We identified the small GTPase, Rac1, as the critical signaling mediator linking TRPML1 to changes in actin polymerization, as blocking Rac1 activity completely mitigated the effect of TRPML1 activation on F-actin. Importantly, utilizing a genetically encoded Ca^2+^ indicator (GCaMP6s), we observed spontaneous TRPML1-derived calcium transients in both OPCs and oligodendrocytes. Collectively, our work has identified a novel function of TRPML1 in regulation of actin dynamics, implicating lysosomal activity in homeostatic regulation of OL maturation and myelination initiation.

Endolysosomes were primarily regarded as the degradative organelle of the cell, with few roles outside of this “trash can” task; however, emerging evidence has positioned them as essential mediators of numerous homeostatic functions, including metabolic signaling^52^, plasma membrane repair^53^, and inter-organelle ion and metabolite exchange^54,55^. Within the oligodendrocyte lineage, less is known about what potentially specialized roles endolysosomes play in regulating both the maturation and myelination process. The transport of the major myelin protein, PLP, is known to be dependent on lysosomal exocytosis^16,18^; furthermore, the myelin proteins, MAG and MOG, and the OPC proteoglycan, NG2, depend on endocytic recycling through the lysosome^56^. Despite this, the localization of the endolysosome throughout the oligodendrocyte maturation process had not been investigated. Our data demonstrate that OPCs undergoing proliferation have few endolysosomes and they primarily reside within the soma (Fig. 1). Once cells begin undergoing differentiation, there is a shift in endolysosome localization into nascent processes where they undergo both anterograde and retrograde trafficking in basal conditions (Fig. 1). Three days after differentiation, when oligodendrocytes are considered mature in our paradigm, there was a significant blunting of endolysosomal motility (Fig. 1). These data suggest that during active differentiation, when developing oligodendrocytes are rapidly changing their transcriptome and need to deliver essential myelin proteins to the plasma membrane, endolysosomes are undergoing not only an increase in their numbers but rapid mobilization into developing processes. It is presently unclear why endolysosomes in mature oligodendrocytes become stationary compared to those at earlier time points. It is possible that at this maturation stage, there is less demand for rapid myelin protein and lipid turnover, which is supported by *ex vivo* data from rodents^57^.

The question arises as to whether these endolysosomes remain inside compacted myelin sheaths to degrade cargo and potentially alter signaling cascades. Previous work investigating autophagosome formation and maturation in MBP+ oligodendrocytes revealed that LysoTracker^+^ puncta were found in branches, which are small channels that may correspond to intramyelinic cytoplasmic channels^57^. These cytoplasm-rich channels are necessary to provide metabolic support, maintain functional axon-glial units, and regulate myelin thickness within active neuronal circuits^58^. Interestingly, 2’,3’-cyclic nucleotide 3’-phosphodiesterase (CNP) binds directly to F-actin within these channels and, together they antagonize MBP-induced compaction to maintain cytoplasmic regions within myelin^58^. However, it is not known how the F-actin within these channels is reorganized as myelin undergoes subtle changes in response to plasticity. It seems plausible that endolysosomes localized within these distal cytoplasmic channels, in addition to degrading myelin lipids and proteins, may also generate calcium nanodomains needed to alter the actin cytoskeleton^14^.

We unexpectedly found that activating or inhibiting TRPML1 had no effect on lineage marker expression in a 2D culture (Fig. 3); this contrasted with our myelinating nanofibers cultures, where inhibition of TRPML1 resulted in significantly fewer PLP+ mature oligodendrocytes (Supplementary Fig. 4). This is the first study, to our knowledge, to test the behavior of OPCs grown in a monolayer versus those grown on nanofiber scaffolds following pharmacologic treatment. It is presently unclear why OPCs grown and differentiated on nanofibers form fewer mature oligodendrocytes following MLSI1 treatment. It is likely that artificial nanofiber system recapitulates more closely *in vivo* conditions, as maturing OPCs need to not only differentiate but also physically contact the fibers and wrap around them to form myelin. Interestingly, this decrease in mature oligodendrocytes does not appear to be driven by the loss of premyelinating oligodendrocytes, a population of terminally differentiated cells that are not OPCs but have not started to form myelin^59,60^, as there were no significant changes in total cell number (DAPI+). Apoptosis of differentiating OPCs that fail to mature into myelin forming oligodendrocytes is common and continues throughout adulthood^61,62^. Thus, based on our data, TRPML1 appears to control the ability of OPCs on nanofibers to differentiate into mature myelinating OLs, which is not recapitulated in 2D monolayer culture.

The most striking phenotype we observed on both 2D monolayer and 3D nanofiber cultures was alterations to oligodendrocyte process morphology, including length and complexity. Activation of TRPML1 at the time differentiation resulted in more complex branch morphology as assessed via Sholl analysis, while inhibition of the channel resulted in significantly stunted process length and complexity (Fig. 4). These changes in oligodendrocyte morphology coincided with measurable changes in F-actin content and polymerization (Fig. 5). The alterations in actin polymerization induced by activation or overexpression of TRPML1 were most prominent during the very early stages of maturation (e.g. 24 hours after switch to DM). This is unsurprising given the massive morphologic alterations that occur at the early stages of transition from an OPC to a premyelinating oligodendrocyte^8^. Previous studies have clearly demonstrated that dynamic actin-cytoskeletal mechanisms underlie essential aspects of OL differentiation, including process outgrowth and branching, and the initiation of myelination. Numerous intracellular mediators of actin dynamics in OLs have been identified, including Arp2/3, WAVE1^5^, Fyn^63^, and FAK^64^; however, this is the first study to implicate an intracellular organelle in the regulation of actin polymerization in the OL lineage. These data add to the growing evidence that lysosomes, outside of their primarily degradative role, can act as an intracellular platform that link biochemical signals to cytoskeletal changes. While previous work has shown that the cytoskeletal, specifically actin, alterations associated with lysosomal signaling has been implicated in cellular migration^26,65^, our work here highlights a previously undescribed role of lysosomal signaling in process outgrowth and morphology. Intriguingly, our data adds to accumulating evidence that cytoskeletal dynamics are potentially regulated independently from transcriptional gene changes associated with differentiation. Conditional knockout of *Arp3c* or inactivation of Fyn tyrosine kinase, for example, have been shown to greatly reduce OL process outgrowth with no changes observed in MBP, O1, and MAG^7,11^. Together with our data, this strongly suggests that at the premyelinating stages, morphological maturation of OLs can be regulated by actin regulators, such as Arp2/3, Fyn, and now TRPML1, that apparently function independently from those regulating transcriptional myelin gene expression.

It is interesting to note that the changes in morphology that we observed in our 2D monolayer culture were also present in the nanofiber myelinating 3D cultures, in contrast to what we observed with lineage marker expression (Supplementary Figs. 4 & 5). While this may seem initially surprising given the necessary switch from actin polymerization to depolymerization for myelination to occur^6,7^, nanofiber cultures do not recapitulate all aspects of myelination. In addition to lacking external signals from neurons to trigger myelination, the myelin that is typically generated on nanofibers rarely undergoes compaction and its membranes are poorly organized^66^. Thus, nanofiber cultures appear to be a reliable model for myelination initiation, which is a process driven primarily by actin polymerization rather than depolymerization^6,7^.

Our morphological data following TRPML1 activation suggested the following question: how does a lysosomal cation channel regulate actin dynamics? Previous reports looking at the protein interaction network of TRPML1 revealed that it interacts with and colocalizes with several Rho GTPases, including Rac1^47^. This is in agreement with other studies that have demonstrated that Rac1, while predominantly found on the plasma membrane, associates with LAMP1+ structures^65,67^. Our studies showed robust activation of Rac1 (Rac1 bound to GTP) as early as 15 minutes after treatment with MLSA1 (Fig. 6a). Previous *in vitro* data demonstrated that Rac1 is a positive regulator of OPC morphological differentiation; however, *in vivo* data from Rac1 mutants under a *Cnp* promoter did not reveal any change in the number of myelinated fibers^68^. While this may initially appear as contradictory data, it is important to note the myelin in Rac1 mutant mice had myelin outfoldings, defined as areas of internodes where the myelin sheath protrudes away from the axon surface^68^. These myelin outfoldings are also found in mice with attenuated calcium signaling (CalEx mice) that also results in perturbation of actin polymerization^14^. Therefore, Rac1 seems poised to be at the nexus between calcium and actin signaling, ultimately impacting both OL morphology and appropriate myelination.

Rac1 sits upstream of several known actin-regulating pathways, including N-WASP, WAVE1, and PAK1^69^. We observed robust phosphorylation (activation) of PAK1 downstream of Rac1 activity in differentiating OLs treated with MLSA1 (Fig. 6b). PAK1, upon its activation by Rac1, phosphorylates Lim kinase (LIMK), which then phosphorylates and inactivates the actin-severing protein Cofilin. Indeed, we saw that Cofilin is phosphorylated, and thus inactivated, in response to TRPML1 activation (Fig. 6c). This coincided with a shift towards actin polymerization, as assessed via the ratio of F/G actin (Fig. 5e). Inactivation of PAK1 activity both *in vitro* and in zebrafish has previously been shown to significantly impact oligodendrocyte morphology and process complexity, similar to what we observe when TRPML1 is inhibited during maturation^37^. While we did not examine other pathways that are downstream of PAK1 activation, this protein is known to regulate other signaling pathways implicated in oligodendrocyte development and myelination, including the AKT and MAP kinase (MAPK) pathways^69^. Additionally, both PI-3K and AKT can directly activate PAK1 in a GTPase-independent manner^70,71^. It remains to be seen whether regulation of PAK1 activity by TRPML1 also influences other downstream pathways like AKT.

As a non-selective lysosomal cation channel, TRPML1 is permeable to Ca^2+^, Fe^2+^, Zn^2+^, Na^+^, and K^+^. Of these cations, calcium is the most likely candidate linking TRPML1 to Rac1 activation. Changes in intracellular calcium are known to regulate the translocation and activation of Rac1^72^. Additionally, in other cell types, calcium signaling plays essential roles in regulating cytoskeletal dynamics, exocytosis, and gene expression, all processes linked to OL maturation and myelination. The role of calcium signaling has been extensively studied in OPCs, where it plays a necessary role in OPC morphology and proliferation; however, less attention has been paid to its role in differentiating and premyelinating OLs. A recent study found that genetic attenuation of calcium did not affect the number of myelinated axons but instead resulted in significant morphological changes to myelin sheath geometry and reduction in actin filaments^14^. The source of the calcium necessary for appropriate actin filament levels and the downstream molecular mediators are unknown. The work presented here demonstrates that calcium efflux through TRPML1 is responsible, at least in part, for the actin filament increase needed for appropriate process growth and complexity. Utilizing a genetically encoded calcium indicator tagged to the amino terminal of TRPML1, we have demonstrated the presence of spontaneous, lysosome-derived calcium transients in OPC and differentiating OL processes (Fig. 2a and Supplementary Video 2). We observed two distinct kinds of calcium transients in OL processes: fast and prolonged. Fast transients appeared and dissipated within a ∼three second period and were generally restricted to small (< 5µm) area of the process (Fig. 2a-b); prolonged calcium lysosomal efflux, on the other hand, occurred over a ∼nine second period. Additionally, these prolonged events typically spanned the entire length of the OL process (Fig. 2a-b). Regardless of if they were fast or prolonged, these calcium events never spilled over into neighboring processes or into the soma, suggesting that these are restrictive calcium microdomains. These microdomains are reminiscent of those found in newly myelinating OLs that are associated with specific myelin remodeling outcomes^73,74^. High-amplitude, long-duration calcium transients preceded myelin sheath retractions, while lower-amplitude, shorter-duration ones positively correlated with sheath growth^73,74^. Therefore, it is possible that highly localized calcium signaling during the pre-myelination stage regulates actin polymerization in a similar manner, where fast transients are associated with process outgrowth, while prolonged ones predict process retraction. Ongoing experiments in our laboratory are currently investigating the relationship between calcium and actin polymerization.

We complemented our *in vitro* work by assessing myelination and mature oligodendrocyte number in a genetic model of MLIV. *Mcoln1^-/-^*mice contain a neo cassette replacing exons 3-4 and part of exon 5 of the *Mcoln1* gene, which abolishes gene expression^75^. Importantly, these mice display all the hallmarks of MLIV, including motor deficits, retinal degeneration, and elevated levels of plasma gastrin^30^. While previous work has characterized potential myelination deficits in these mice^31,50,76^, no work thus far has investigated a potential mechanism underlying these observations. As previously reported in P10 mutant mice, we observed a significant reduction in PLP staining intensity in both the corpus callosum and motor cortex in P21 *Mcoln1^-/-^*mice (Fig. 8). These data suggest that both gray and white matter derived OLs, despite reported differences in proliferation rates, response to injury, and myelination capacity^77–79^ may be equally susceptible to loss of TRPML1 and highlights the fundamental role this protein plays in regulating developmental myelination. Interestingly, we observed a slight, but significant, increase in PLP immunoreactivity in animals containing one *Mcoln1* allele. Previous work with these mice did not investigate whether animals with one functional allele exhibit any alterations compared to wild-type mice. Genetic compensation, whereby either related genes or the functional allele are upregulated, has been proposed to occur in response to deleterious mutations but not after translational or transcriptional knockdown^80,81^. While we have not investigated expression levels of TRPML1 or its related proteins, TRPML2 and TRPML3, in *Mcoln1^+/-^* mice, it is possible that degradation of mutant mRNA triggers transcriptional adaptation in these animals. Despite changes in myelination, we did not observe drastic alterations in mature oligodendrocyte number (ASPA+); instead, significant differences were only seen in the corpus callosum and superficial layers of the motor cortex (Fig. 8). These data are in line with our *in vitro* findings, where we did not observe any changes in oligodendrocyte lineage marker expression but saw massive morphological alterations when TRPML1 was either activated or inhibited.

As we observed in our studies using primary rat oligodendrocytes, *Mcoln1^-/-^* mice displayed a significant reduction in levels of pPAK1 within Olig2+ cells compared to wild-type and *Mcoln1^+/-^*animals (Fig. 9). These data suggest that the mechanisms we elucidated *in vitro* are potentially involved in the myelination deficits present within these mice. It was recently shown that AAV-mediated CNS-targeted gene transfer of the human *MCOLN1* in the MLIV mouse model rescued motor deficits and improved myelination within the corpus callosum^82^. It would be interesting to see if the observed restoration in myelination is associated with normalization of PAK1 activity within the OL lineage.

In summary, we discovered that the lysosomal cation channel TRPML1 plays an essential role in regulating oligodendrocyte actin polymerization and morphology through Rac1/PAK1 activity. Furthermore, our work suggests that TRPML1 activity may serve as a mediator between lysosomal membrane signaling, via generation of the endogenous agonist of TRPML1, PI(3,5)P_2_, and the Rac1/PAK pathway to modulate oligodendrocyte actin organization. These data add to a growing body of evidence that demonstrate a role for the lysosome in regulating critical homeostatic functions within different cell types and highlights how examining lysosomes outside of only gene expression may reveal novel phenotypes. Additionally, this work has implications not only for hypomyelination observed in MLIV patients, but also other diseases where white matter impairment and dysfunctional TRPML1 have been implicated, including Alzheimer’s disease and multiple sclerosis.

## Methods Animals

All procedures involving animals were approved by the Institutional Animal Care and Use Committee (IACUC) of the Children’s Hospital of Philadelphia. Animals were group-housed in a CHOP animal facility with a 12:12 h light/dark cycle. Mice were given ad libitum access to food and water. *Mcoln1^-/-^* frozen embryos were purchased from Jackson Laboratories (JAX stock #027110) and rederived at the CHOP Transgenic Core. *Mcoln1^-/-^* mice contain a neo cassette that replaces exons 3-4 and part of exon 5 of the *Mcoln1* gene, which abolishes gene expression (Venugopal et al 2007). The *Mcoln1* line was maintained by crossing *Mcoln1^+/+^*and *Mcoln1^+/-^* progeny. Since homozygous animals display hind-limb paralysis by 6.5 months and have a life expectancy of 8 months, we utilized animals from all genotypes at 3 months of age or younger. The following primers were used for genotyping: Common *Mcoln1* reverse 5’-primer CAGTGTGAGGTTCTTCTAACTGG; WT forward 5’-primer GGGAGTTAAACAGTGAAGAAGG; Mutant forward 5’-primer CAGCTGGGGCTCGACTAGA. All resulting progeny from these crosses were used. Sprague-Dawley rats for primary OPC cultures were purchased from Charles River Laboratories. Male and female mice were used for *ex vivo* experiments. For cell culture studies, brains of both sexes were pooled together to obtain sufficient cell numbers.

## Primary rat oligodendrocyte precursor cell cultures

Primary rat glial cells were isolated from brains (2-4 brains per biological replicate) of male and female postnatal day 1 Sprague-Dawley rats (Charles River Laboratories) and plated on T75 flasks^83^. Upon reaching confluency, OPCs were purified using the “shake-off” method^84^. Briefly, T75 flasks were rotated on an orbital shaker set to 250 rpm and incubated overnight at 37C. The following day, cells were filtered using a 20-µm nylon net (Merck Millipore), followed by centrifugation at 450 *g* for 5 min at 4C. The supernatant was discarded, and the pellet resuspended in 5 mL Neurobasal medium (Life Technologies) supplemented with B27 and glutamine and incubated on a bacteriological Petri dish for 15 mins at 37°C. The supernatant was collected and centrifuged at 450 *g* for 5 min at 4C. The pellet was resuspended in Neurobasal medium supplemented with B27, glutamine, and growth factors: PDGF (2 ng/mL), NT3 (1 ng/mL), and FGF (10 ng/mL), and plated on 24-well plates with coverslips (immunocytochemistry), 35 mm dishes with glass coverslips (live cell imaging), 10 cm dishes (immunoblotting), and 12-well cell culture inserts containing 700 nm aligned nanofibers (nanofiber culture; Nanofiber Solutions). Once OPCs reached ∼60-70% confluency, growth medium was replaced with differentiation medium containing 50% DMEM, 50% Ham’s F12, pen/strep, 2mM glutamine, 50 ug/mL transferrin, 5 ug/mL putrescine, 3 ng/mL progesterone, 2.6 ng/mL selenium, 12.5 ug/mL insulin, 0.5 ug/mL T4, 0.3% glucose, and 10 ng/mL biotin. At the time of differentiation, cells were treated with vehicle (DMSO), MLSA1 (20 μM), MLSI1 (15 μM), or NSC23766 (100 μM) for 24-72 h prior to experimental end points. On average, typically 30-50% of OPC differentiation into mature oligodendrocytes is observed in untreated conditions.

## Reagents

Reagents and chemicals used were LysoTracker Green DND-26 (Thermo Fisher Scientific, #L7526), LysoTracker Red DND-99 (Thermo Fisher Scientific, #L7528), MLSA1 (Millipore Sigma, #SML0627), MLSI1 (MedChemExpress, #HY-134818), Phalloidin Alexa Fluor 594 (Thermo Fisher Scientific, A12381), latrunculin A (Millipore Sigma, #428021), jasplakinolide (Millipore Sigma, #J4580), NSC23766 (Tocris Bioscience, #2161), TPC2-A1-N (MedChemExpress, #HY-131614), SG-094 (MedChemExpress, #HY-148816),

## Antibodies

The following antibodies were used for immunohistochemical analysis of brain tissue sections: aspartoacylase (ASPA; GeneTex, GTX110699; 1:500), proteolipid protein^85^ (PLP; AA3 rat hybridoma; 1:2), Olig2 (Abcam ab109186; 1:200), and pPAK1 (Thr423, Invitrogen PA5-12844; 1:25). The following antibodies were used for immunocytochemistry *in vitro*: A2B5^86^ (OPCs; mouse IgM hybridoma; 1:2), galactocerebroside^87^ (GalC; immature/mature oligodendrocytes; mouse IgG3 hybridoma; 1:5), proteolipid protein^85^ (PLP; mature oligodendrocytes; AA3 rat hybridoma; 1:2). The following antibodies were utilized for immunoblotting: pPAK1 (Thr423)/pPAK2 (Thr402) (Cell Signaling Technology, #2601; 1:1000), PAK (Cell Signaling Technology, #2602; 1:1000), pCofilin (Ser3) (Cell Signaling Technology, #3311; 1:1000), Cofilin (Cell Signaling Technology, #5175; 1:1000), Rac1 (Cell Biolabs, included in #STA-401-1; 1:1000), beta actin (β-actin; Sigma-Aldrich, #2066; 1:6000), and alpha tubulin (α-tubulin; Sigma-Aldrich, #T5168; 1:10000).

## Immunoblotting

Whole cell extracts of primary rat oligodendrocytes were prepared in ice-cold lysis buffer (25 mM Tris (pH 7.4), 10 mM EDTA, 10% SDS, 1% Triton-X 100, 150 mM NaCl, protease and phosphatase inhibitor cocktail) followed by sonication and centrifugation at 10,500 *g* at 4C for 30 min. Protein concentrations were determined using bicinchoninic acid (BCA) assay following the instructions of the manufacturer (Pierce). Protein (7.5-20 ug) were loaded onto a 4-12% Bis-Tris gradient gel and after separation, proteins were transferred to Immobilon-FL membranes. Primary antibodies were incubated overnight at 4C with primary antibodies in 2X NAP. The following day, membranes were incubated with fluorescent probe-conjugated secondary antibodies in TBS-T. Membranes were visualized using an Odyssey Infrared Imaging System (LiCOR).

## Rac1 activation assay

The Rac1 activation assay was performed using Cell Biolabs Rac1 Activity Assay Kit (#STA-401-1). Briefly, cells were lysed as described above and the active form of Rac1 (GTP-Rac1) was selectively pulled down from the lysate with the p21-binding domain (PBD) of PAK agarose beads. Subsequently, the precipitated GTP-Rac1 was detected by immunoblot analysis as described previously. Each assay also consisted of lysates that were loaded with either GTPγS or GDP as positive or negative controls, respectively. A separate immunoblot was run to evaluate total levels of Rac1 in each lysate.

## F/G actin ratio

F/G actin ratio was assessed as previously described. Briefly, cells were lysed in cold lysis [10 mM K_2_PO_4_, 100 mM NaF, 50 mM KCL, 2 mM MgCl_2_, 1 mM EGTA, 0.2 mM DTT, 0.5% Triton-X 100, 1 mM sucrose (pH 7.0)], and centrifuged at 15,000 x *g* for 30 min. Separation of F-actin and G-actin was achieved in that F-actin is insoluble (pellet) in this buffer, whereas G-actin is soluble (supernatant). The G-actin supernatant was transferred to a fresh tube, and the F-actin pellet was resuspended in lysis buffer plus an equal volume of a second buffer [1.5 mM guanidine hydrocholoride, 1 mM sodium acetate, 1 mM CaCl_2_, 1 mM ATP, 20 mM tris-HCl (pH 7.5)] and then incubated on ice for one hour with gentle mixing every 15 min to convert F-actin into soluble G-actin. Samples were centrifuged at 15,000 x *g* for 30 min and the supernatant (containing F-actin converted to G-actin) was transferred to a fresh tube. F-actin and G-actin samples were loaded with equal volumes and analyzed via immunoblot. Latrunculin A (5 μM, 2 hr), a potent actin polymerization inhibitor, and jasplakinolide (5 μM, 2 hr), an inducer of actin polymerization, were used as internal controls for the assay.

## Infection of primary OPCs

One to two days after isolation, primary OPCs were infected with one of the following lentiviral constructs: pLV-Olig2-GCaMP6s-rMcoln1 (MOI: 10), pLV-CMV-eGFP (MOI: 10), pLV-eGFP-Olig2-rMcoln1 shRNA1-3 (MOI: 10), or pLV-eGFP-Olig2-scramble (MOI: 10). 24 hours after infection, virus-containing media was removed and either fresh media with growth factors or differentiation media depending on the experiment. Infection efficiency ranged from 21.4-32.9%.

## Immunostaining of primary oligodendrocytes

At the specified day of differentiation, coverslips were washed three times in DMEM/F12 and incubated in primary antibodies to surface antigens (A2B5 and GalC) for 15 min at RT. Coverslips were then rinsed three times with DMEM/F12 and incubated in appropriate fluorescent-conjugated secondary antibodies for 15 min at RT. Cells were then rinsed three times with DMEM/F12, fixed with 4% paraformaldehyde for 10 min, and permeabilized with 0.1% Triton-X-100 for 5 min. Cells were then incubated with either primary antibodies for internal antigens (PLP) for 15 min or phalloidin-Alexa Fluor 594 for 1 hr at RT. Coverslips were rinsed three times and incubated with appropriate secondary antibodies for 15 min at RT. Lastly, cells were rinsed, counterstained with DAPI for 5 min, and then mounted on slides with ProLong Gold antifade reagent.

Cells were imaged either using a Keyence BZ-X-700 digital fluorescent microscope (Keyence Corporation) for OPC and oligodendrocyte cell counts or a DMi8 Leica inverted confocal microscope equipped with a 40x/1.3 NA objective. Quantification of oligodendrocyte lineage counts were conducted by hand counting the number of OPCs, immature oligodendrocytes, mature oligodendrocytes, and DAPI+ cells across 25 fields/coverslips and three coverslips per condition for each biological replicate. The percentage of each cell type was calculated and normalized to the untreated condition. For phalloidin corrected total cell fluorescence (CTCF), imaging fields were chosen at random and Z-stacks from five fields/coverslip and three coverslips/condition for each biological replicate were acquired. ROIs were individually drawn around each cell in an image and integrated density was measured using Fiji. Additionally, three to five background measurements were taken and CTCF was calculated per cell using the formula (Integrated Density - (Area of selected cell x Mean fluorescence of background readings).

## Lysosome localization imaging

Purified OPCs and differentiating oligodendrocytes were loaded with LysoTracker Green DND-26 (75 μM) for 30 minutes prior to being washed and placed in Hibernate E supplemented with 2% B27 and 2% L-glutamine (HibE imaging solution) for imaging. Live-cell imaging was performed using an Olympus Fluoview i10 with a humidified, temperature-controlled microscope enclosure and equipped with a 60x/1.2 NA water immersion objective. The entire well was imaged using phase contrast microscopy and five random areas were selected from this image. Images were acquired every ten seconds in the 488 channel over a 15 min period. Kymographs of selected OPC and oligodendrocyte processes were generated from primary processes over a 50 μm span using the KymoToolBox Fiji plugin and a line width of 5 pixels. To measure vesicle dynamics, individual tracks were manually traced using the segmented line tool. For each vesicle tracked, the KymoToolBox plugin calculated the total distanced traveled, the percentage of time moving in an anterograde, retrograde, or stalled manner, and the mean speed (total distance traveled divided by total time tracked).

## Calcium imaging

OPCs and oligodendrocytes expressing the GCaMP6s-TRPML1 construct were imaged on a BioVision spinning disk confocal microscope system with a Leica DMi8 inverted widefield microscope, a Yokagawa W1 spinning disk confocal, and a Photometrics Prime 95B scientific complementary metal-oxide-semiconductor camera. Images were acquired with the VisiView software using a 63x/1.4 NA oil immersion objective. For images derived from the same biological replicate, the same acquisition parameters were used across all conditions. The microscope is equipped with an adaptive focus control to maintain a constant focal plane during live-cell imaging and an environmental chamber to maintain 37°C. Images were acquired every second over a two min period. Calcium imaging data were analyzed using Fiji to generate ROIs around the somas and processes of OPCs and oligodendrocytes and rate and amplitude measurements were extracted.

## Immunohistochemistry

Mice were anesthetized with isoflurane mixed with oxygen (∼3% flow rate with 1.0 L/min oxygen) and were intracardially perfused first with ice-cold PBS and then 4% paraformaldehyde (pH 7.4). Brains were removed from the skull, and the olfactory bulb and cerebellum were removed using a brain matrix. Brains were post-fixed in 4% paraformaldehyde for 4h followed by immersion in 30% sucrose solution for at least 24h. Brains were cryopreserved and embedded in optimal cutting temperature compound (OCT) followed by coronal sectioning (20 µm) on a cryostat (Leica Microsystems). For PLP staining, sections were immersed in 100% ethanol for 10 min at room temperature and blocked with a blocking solution containing 10% normal goat serum, 2% bovine serum albumin, and 0.1 Triton-X in 1X PBS for 30 minutes. The sections were then incubated overnight at 4°C with PLP (hybridoma) diluted in the blocking solution (1:2). The next day, sections were rinsed three times in 1X PBS and incubated with fluorescent secondary antibodies in the blocking solution (1:200) for 1 h at RT. Sections were rinsed three times in 1X PBS before being incubated overnight at 4°C with ASPA (GeneTex, GTX110699) in a 2%NGS solution (1:500). The next day, sections were rinsed three times in 1X PBS and incubated with fluorescent secondary antibodies in a 2% NGS solution (1:200) for 1 h at RT. Sections were rinsed three times in 1X PBS prior to being counter-stained with DAPI for 5 min and mounted in ProLong Gold anti-fade reagent. Images of the entire section were acquired on a Keyence BZ-X-700 digital fluorescent microscope and stitched together using BZ-X-700 Analyzer software. To quantify PLP integrated density, nine images across the corpus callosum and motor cortex from each animal were analyzed. The channel of interest was separated and thresholded to isolate staining from background, and integrated density was calculated using ImageJ. Quantification of cell number (ASPA and DAPI) was performed in a similar manner and the number of ASPA+ and DAPI+ cells were hand counted using ImageJ.

## Tyramide stain and amplification

Serial sections from the same animals used for PLP and ASPA quantification were used to sequential stain for pPAK1 and the oligodendrocyte lineage marker Olig2. Antigen retrieval was performed using L.A.B. (liberate antibody binding) solution for 20 min at RT followed by quenching of endogenous peroxidase activity with hydrogen peroxide and methanol for 30 min at RT. Tissue was blocked with 10% normal goat serum for 1 h at RT before incubation with primary overnight at 4°C in 2% normal goat serum. Secondary antibody (goat anti-rabbit poly-HRP) incubation was performed for 1 h at RT, then amplified with Alex Fluor conjugated tyramide (Invitrogen, 488 for pPAK1). Tissue was stained for Olig2 using the above-mentioned immunohistochemistry protocol. Tissue was counter-stained with DAPI and mounted in ProLong Gold anti-fade reagent. Images were acquired on a DMi8 Leica inverted confocal microscope using a 40x objective/1.3 NA. Integrated density of pPAK within each individual Olig2+ oligodendrocyte lineage cell was measured as outlined in the previous section. Five images were acquired per animal and then averaged into a single data point per animal.

## Statistics and analysis

All in vitro experiments were performed with at least *n*= 3 biological replicates, and each biological replicate used OPCs derived from a different litter of P0-P4 rat pups. Outliers were neither encountered not removed from any analysis. Experimenters were blinded to treatment conditions or genotype when performing analyses. Data are represented as mean ± SEM for *in vitro* and *in vivo* experiments. For all experiments, statistics were performed on the means from biological replicates (cells from different preps or animals); each mean was calculated from the technical replicates within a biological replicate. This distribution of data is represented using Superplots^88^; each smaller dot represents a value from a single cell or image, while larger dots represent means from biological replicates.

## Acknowledgements

This project was supported by the following grants: National Multiple Sclerosis Society TA-2204-39435 and TA-2507-44763 (LKF), NIH R01 MH098742 (JBG and KJS) and NIH R01 MH126773 (JBG and KJS). We thank the laboratory of Sandra Maday, PhD at the University of Pennsylvania for use of their BioVision spinning disk confocal microscope. We thank the laboratory of Michael Robinson, PhD at the Children’s Hospital of Philadelphia for the use of the Odyssey Infrared Imaging System and the Leica DMi8 confocal microscope. We thank the CHOP Transgenic Core for rederiving the *Mcoln1^-/-^* mouse line. The authors declare no competing financial interests.

