## Supplemental Figure Legends for "The lysosomal cation channel TRPML1 regulates the oligodendrocyte cytoskeleton"

**Supplementary Figure Legends**

**Supplementary Figure 1. TRPML1 modulates lysosomal motility in early differentiating oligodendrocytes. (a)** Endolysosome mean speed significantly increased upon treatment with MLSA1 in 24-differentiated OLs; conversely, inhibition of TRPML1 with MLSI1 drastically blunted mean speed of endolysosomes. (**b)** As observed with endolysosome speed, the cumulative distance traveled of individual endolysosomes were altered with activation or inhibition of TRPML in 24-hour differentiated OLs. (**c)** Similar to 24-hour differentiated OLs, activation or inhibition of TRPML1 within OLs differentiated for 48 hours significantly altered endolysosome mean speed, with activation of TRPML1 increasing mean speed and MLSI1 decreasing it. (**d)** In 48-hour differentiated OLs, endolysosome cumulative distance was significantly modulated by activating or inhibiting TRPML1. (**e)** Activation of TRPML1 significantly reduced the percentage of endolysosomes that were classified as “stationary” compared to control and MLSI1-treated OLs that had been placed in differentiation medium for 24 hours. (**f)** MLSA1 treatment resulted in a slight decrease in “stationary” endolysosomes in 48-differentiated OLs. N=3 biological replicates (preps); small symbols represent individual cells; each shape and color are from a different biological replicate. One-way ANOVA followed by Tukey’s post-hoc test (a-d).

**Supplementary Figure 2. The genetically-encoded calcium indicator, GCaMP6s-TRPML1 localizes to endolysosomes in OPCs and oligodendrocytes.** Representative images of GCaMP6s-TRPML1 and the endolysosome dye, Lysotracker Red, in lentiviral-infected OPCs. There is extensive overlap of their fluorescence signals, demonstrating appropriate localization of the GCaMP6s-TRPML1 construct; scale bar = 10 μm.

**Supplementary Figure 3. Knockout or overexpression of TRPML1 does not affect oligodendrocyte lineage marker expression. (a)** Representative images of three-day differentiated OL cultures infected with scramble, *Mcoln1* shRNA, or *Mcoln1*-overexpression lentiviral constructs and stained for A2B5, GalC, PLP, and DAPI. Scale bar = 50 µm. **(b)** No changes were observed in the percentage of cells expressing the OPC marker, A2B5, across all groups. **(c)** There were no significant alterations in the percentage of cells expressing the immature/mature oligodendrocyte marker, GalC, across all groups. **(d)** As observed with A2B5 and GalC, there were no significant differences in the percentage of cells labeled with the mature oligodendrocyte marker, PLP, across all groups. **(e)** Total cell number, as measured by DAPI+ nuclei staining, was not significantly different across any of the groups. N=3 biological replicates (preps). One-way ANOVA followed by Tukey’s post-hoc test; all p>0.05.

**Supplementary Figure 4. Activation or inhibition of TRPML1 in myelinating OL cultures affects oligodendrocyte lineage marker expression. (a)** Representative images of three-day differentiated OL cultures placed on 700 nm artificial nanofibers and treated with vehicle, MLSA1, or MLSI1. Scale bar = 10 µm. **(b)** Inhibition of TRPML1 resulted in a significant increase in the percentage of cells expressing the OPC marker A2B5. **(c)** Treatment with MLSI1 caused a drastic reduction in the percentage of PLP+ cells on artificial nanofibers. **(d)** There were no observed changes in total cell number (DAPI+) in any of the treatment groups. N=3 biological replicates (preps). One-way ANOVA followed by Tukey’s post-hoc test.

**Supplementary Figure 5. TRPML1 controls oligodendrocyte morphology during early myelination. (a)** Representative images of three-day differentiated OL cultures plated on 700 nm artificial nanofibers, treated at the time of differentiation with vehicle, MLSA1, or MLSI1, and stained with A2B5, PLP, and DAPI. Scale bar = 10 µm. **(b)** MLSA1-treated PLP+ cells exhibited significantly greater morphologic complexity as measured via Sholl analysis. Conversely, MLSI1-exposed cells had simpler morphology, particularly at distal regions of the processes. **(c)** Quantification of Sholl analysis by area under the curve. **(d)** Primary process length (processes originating from the soma) was significantly decreased in MLSI1-treated cultures. N=3 biological replicates (preps). Two-way ANOVA or one-way ANOVA followed by Tukey’s post-hoc test; *p<0.05, **p<0.01, ***p<0.001**.**

**Supplementary Figure 6. Knockdown or overexpression of TRPML1 alters actin filament content in early differentiating oligodendrocytes. (a)** Representative images of one-day differentiated OL cultures infected with scramble, *Mcoln1* shRNA, or *Mcoln1* overexpression lentiviral constructs and stained with phalloidin. Scale bar = 10 µm **(b)** Knockdown of TRPML1 significantly decreases F-actin content in differentiating OLs, while overexpression results in an upregulation of actin filaments. N=3 biological replicates (preps); small symbols represent individual cells; each shape and color are from a different biological replicate. One-way ANOVA followed by Tukey’s post-hoc test.

**Supplementary Figure 7. Activation or inhibition of TPC2, a Ca^2+^-permeable lysosomal channel, does not influence F-actin content in differentiating oligodendrocyte cultures. (a)** Representative images of one-day differentiated OL cultures treated with MLSA1, TPC2-A1-N (TPC2 agonist; 10 µM), or SG-094 (TPC2 antagonist; 10 µM) and stained with phalloidin; scale bar = 10 μm. **(b)** As previously demonstrated, activation of TRPML1 increased F-actin content in one-day differentiated OL cultures, while MLSI1-treated cultures significantly reduced phalloidin staining intensity; on the other hand, activation or inhibition of TPC2 had no impact on F-actin content. **(c)** Similar results were observed in two-day differentiated OL cultures. N=3 biological replicates (preps); small symbols represent individual cells; each shape and color are from a different biological replicate. One-way ANOVA followed by Tukey’s post-hoc test.

**Supplementary Figure 8. Inhibition of Rac1 does not affect OL lineage marker expression. (a)** Representative images of three-day differentiated OL cultures treated with vehicle, MLSA1, NSC23766 (Rac1 inhibitor; 50 µM), or NSC23766+MLSA1 and stained for A2B5, GalC, PLP, and DAPI; scale bar = 50 μm. **(b)** No changes were observed in the percentage of cells expressing the OPC marker, A2B5, across all treatment groups. **(c)** There were no significant alterations in the percentage of cells expressing the immature/mature oligodendrocyte marker, GalC, across all groups. **(d)** As observed with A2B5 and GalC, there were no significant differences in the percentage of cells labeled with the mature oligodendrocyte marker, PLP, across all treatment groups. **(e)** Total cell number, as measured by DAPI+ nuclei staining, was not significantly different across any of the groups. N=3 biological replicates (preps). One-way ANOVA followed by Tukey’s post-hoc test; all p>0.05.

**Supplementary Figure 9. *Mcoln1^-/-^* mice have impaired myelination in layers I-III of the motor cortex. (a)** TRPML1 KO mice have a significant reduction in PLP staining intensity compared to wild-type and heterozygous littermates. **(b)** There were no significant differences observed in the percentage of ASPA+ cells across all genotypes. N=5 animals per genotype; small symbols represent individual sections. One-way ANOVA followed by Tukey’s post-hoc test.
